# Arthropod communities associated with gall-inducing *Aciurina bigeloviae* and *Aciurina trixa* (Diptera: Tephritidae) in New Mexico

**DOI:** 10.1101/2023.09.05.555642

**Authors:** Quinlyn Baine, Emily E. Casares, Daniel W.W. Hughes, Vincent G. Martinson, Ellen O. Martinson

## Abstract

Insect-induced galls are novel structures that serve as habitat to whole communities of associate arthropods that include predators, parasitoids and inquilines. Galling insects are generally under-described, but their associate communities, which can include many specialist organisms, are virtually unknown, particularly in the southwest United States. *Aciurina bigeloviae* (Cockerell 1890) and *Aciurina trixa* Curran 1932 are unusually common and abundant galling flies in New Mexico. The 2 species are sister and occur in sympatric areas but have distinct gall morphologies. We reared all arthropods from 3800 galls from 14 sites in the northern and central regions of the state and as a result characterized the complete communities of both species, including barcode sequences and eclosion phenology. We also investigate interactions of *A. trixa* galls with the abundant inquiline weevil *Anthonomus cycliferus* Fall 1913 and find no measurable effect of inquiline abundance on the size of the emerged adult fly or gall. The total species count is 24 and includes 6 guilds; both *A. bigeloviae* and *A. trixa* communities are richer and more complex than other documented Tephritidae-Asteraceae galling systems. This study highlights the potential of galling insects as ecosystem engineers to maintain large, rich and multi-trophic communities.

## Introduction

Galls are abnormal growths on a plant that are induced by another organism (Redfern 2011). Insect-induced galls are among the most complex, displaying structures and pigments not typically seen on the host plant (Stone and Schönrogge 2003). The gall is considered an extended phenotype of the insect, in that though the structure is made with plant genes and proteins, the morphology and phenology of the gall is determined by the insect (Bailey et al. 2009). A gall is a multi-tissue organ with interior nutritive tissue and exterior defense tissue that provides nutrition and protection to the gall inducer’s offspring (Price et al. 1987, Martinson et al. 2022).

Gall-inducing insects act as ecosystem engineers by creating novel structures that serve as habitats not only for themselves, but also for diverse arthropod functional groups, such as parasitoids, predators and inquilines (Stone and Schönrogge 2003, Bailey et al. 2009, Cornelissen et al. 2016, Luz and Mendonça 2019). Organisms that create habitat for other taxa, ecosystem engineers, play central roles within food webs. In the case of gall-inducers, they create and maintain both trophic and non-trophic resources (i.e., galled plant tissues are potential food, gall structures are potential shelter) (Sanders et al. 2014, Cornelissen et al. 2016) introducing novel structures and new niches to the ecosystem (Price et al. 1987, Cornelissen et al. 2016, Barbosa et al. 2019). Therefore, ecosystems containing gall inducers have the potential to host richer communities than ecosystems with only non-galling herbivore insects (Price et al. 1987). Furthermore, we find that gall-associated arthropods are extremely diverse, not only taxonomically, but functionally (Sheikh et al. 2022, Ward et al. 2022), and in this study show that even when the primary inducers are similar, each inducer hosts distinct associates and increases overall ecosystem biodiversity.

We examined the communities of sister fly species *Aciurina bigeloviae* (Cockerell 1890) and *Aciurina trixa* Curran 1932 (Diptera: Tephritidae) (Fig. 1B-C) that induce galls on *Ericameria nauseosa* (Pall. ex Pursh) G.L.Nesom & G.I.Baird. The species are very similar with almost identical life cycles and similar body morphology, and were briefly synonymized in the taxonomic literature (Steyskal 1984). However, extensive study in sympatric regions of New Mexico found consistent distinction in gall morphology and host plant *variety* specificity, as well as sufficient evidence of reproductive isolation to maintain species designation (Dodson and George 1986). Although *Aciurina* galls are common and abundant throughout the Intermountain West, the power of these species as ecosystem engineers is mostly unknown and the associate communities are almost completely undescribed. Surprisingly, only a single community has been partially characterized for *A. trixa* in Idaho and no communities have been formally characterized for *A. bigeloviae* (Wangberg 1981).

**Figure 1.**
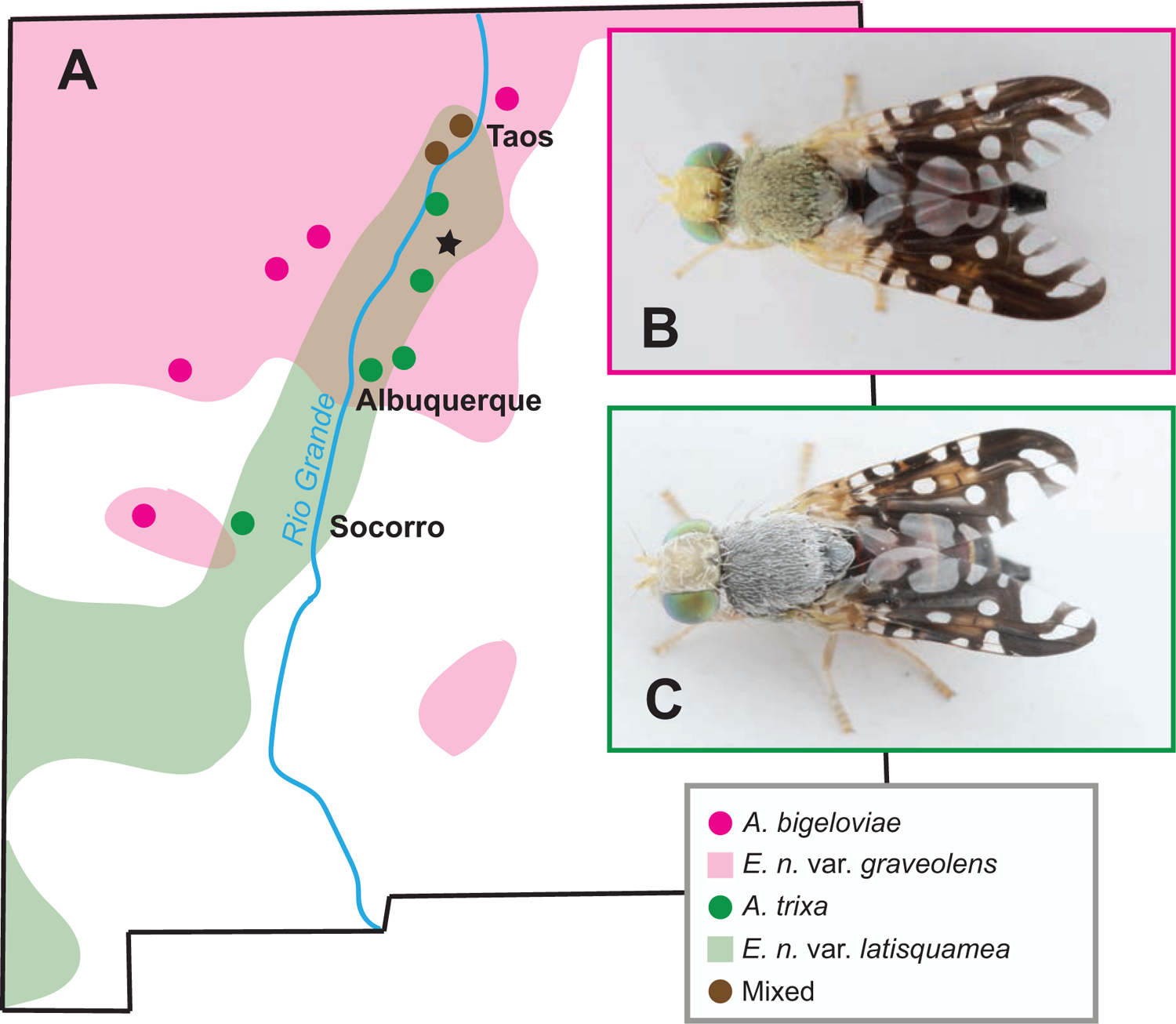
A. Map of New Mexico with gall population collection localities and host plant variety ranges adapted from McArthur 1984. B. *Aciurina bigeloviae* adult female and C. *Aciurina trixa* adult female.

*Aciurina bigeloviae* and *A. trixa* are both univoltine flies that insert single eggs into the base of a leaf bud in spring. The external gall begins to form in late summer and by late fall the developing larva tunnels a single chamber, where it feeds on the lining of nutritional cells at the center of the fully-formed gall. Mature larvae overwinter inside the gall,pupate Mar-May, and subsequently eclose as adults in May-June. In New Mexico, *A. bigeloviae* is specific to *E. nauseosa ssp. nauseosa var. graveolens,* which occurs broadly throughout high-elevation grassland of the northern half of the state, while *A. trixa* is specific to *E. nauseosa ssp. nauseosa var. latisquamea,* which is found in rocky washes along portions of the Rio Grande and Gila River basins. The 2 host plants co-occur in the Rio Grande valley between Santa Fe and Taos, NM and in this region *A. bigeloviae* and *A. trixa* are sympatric and present in high density populations (McArthur 1984, Dodson and George 1986) (Fig. 1A), providing excellent opportunities for comparison of associate communities.

Analysis of gall community structure is essential to our understanding of how insect biodiversity is generated, and therefore how it can be best preserved (Stone and Schönrogge 2003, Feder and Forbes 2010, Hardy and Cook 2010, Brodersen et al. 2018). Enemies can be highly abundant in galling systems and fill many niches (Redfern 2011, Luz and Mendonça 2019). Therefore gall-associated communities are not just taxonomically diverse, but are ecologically complex and represent rare interspecies interactions that are generally severely under-described (Forbes et al. 2016, Weinersmith et al. 2017, Ward et al. 2022). In this paper, we provide a full characterization of species associated with gall-inducers *A. bigeloviae* and *A. trixa* in New Mexico, including host preference, life history, and phenology.

## Methods

### Sample Collection and Rearing

Galls for this study were collected from 14 dense populations in northern and central New Mexico in 2021 and 2022 (Fig. 1A). Each population was sampled for 200 galls across 5-10 highly-galled plants for a total of 3800 galls. Galls were collected in the field on a 10-20 cm length of stem, placed in plastic bags with water at the base, and transported in a cooler to the University of New Mexico (UNM). In 2021, all galls were placed individually in 25.12 mm polystyrene *Drosophila* vials (Fisherbrand) and plugged using narrow mitty plugs (Jaece Industries). In 2022, galls from the first five plants were placed singly into vials, but those from the second five plants were grouped into half-pint jars fitted with a funnel and vial contraption designed to allow emerged insects to fly up and out of the jar, into the vial. All galls were stored during rearing in a plastic tent at a relative humidity of ∼45% and ambient indoor temperature. All rearing containers were checked twice a week for emerged insects, which were then recorded, placed in 100% ethanol, and stored at −20°C. Rearing ceased in August in 2021 and in December in 2022. At the completion of 2022 rearing, all galls without an inducer or primary parasitoid emergence were dissected to determine the cause of inducer mortality. Several additional gall collections were made in the early spring of 2022 that were dissected to examine the life histories of associates.

### Species characterization

Adult insects that occurred in abundance of >1 were identified using the keys cited for each taxon. A subset of emerged insects as representatives of each taxon were accessioned to the Museum of Southwestern Biology (MSB). Habitus photographs were taken on an EOS 40D camera fitted with a 65 mm MP-E macro photo lens (Canon) and mounted on Stackshot macro rail with controller (Cognisys), and then stacked with Zerene Stacker software.

Representative adult specimens and immature specimens removed from early and late season gall dissections were barcode-sequenced to confirm their identity. Whole body DNA was extracted using the DNeasy Blood and Tissue Kit (QIAGEN) with the additional step of homogenization via pestle-grinding. We amplified the mitochondrial *cytochrome oxidase subunit I* (COI) gene using Taq DNA Polymerase (New England BioLabs). Because of inconsistent binding in Chalcidoidea and Ichneumonidae samples, we used four COI primer pairs (LCO1490 and HCO2198 (Folmer et al. 1994); COI pF2 (Simon et al. 1994) and COI 2437d (Kaartinen et al. 2010); BeeCox1F1 and BeeCoxR1 (Bleidorn and Henze 2021); C1-J-1718 and C1-N-2191 (Simon et al. 1994)) and selected the cleanest sequence generated for each species. Amplified products were purified using Exonuclease I (New England BioLabs) and rAPid Alkaline Phosphatase (Roche Diagnostics) using the following thermocycler conditions: 15 min at 37°C, 15 min at 80°C, hold at 10°C. Cleaned amplicons were then submitted for Sanger sequencing by Eurofins Genomics (Louisville, KY). Sequences were trimmed and edited using Geneious Prime 2021.2, then compared with available sequence data from NCBI GenBank and the Barcode of Life Database (BOLD) to determine percent nucleotide similarity to known organisms. Genetic sequences are deposited in BOLD (ACCOM007-23.COI-5P-ACCOM022-23.COI-5P) and Genbank (OR336222-34, OR438293-95).

To make an estimation of the impact of inquiline weevil presence on host *Aciurina* success, we compared the inquiline load (number of individual weevils successfully emerged per gall) within one high abundance population to both the size of each gall and size of adult host fly emerged from each gall. At the time of collection, gall diameter was measured with a Mitotuyo 500-752-20 digital caliper. For fly body size, we used wing length as a proxy. Both wings of each fly were removed at the tegulae and mounted on slides with Euparol medium (Hempstead Halide), then slides were photographed with an Axiocam 208 camera mounted on a Stemi 508 microscope (Zeiss) and measured in ZEN 3.6 (Zeiss). Paired wings were averaged per individual. To determine correlation, we performed linear regression using “lm” in R version 4.1.2 package *stats* (R Core Team 2021) with a poisson distribution.

## Results

Twenty-two species total were identified as gall associates in the aggregate community including 11 species shared between the 2 hosts (Table 1). Nine exclusive and 20 total species are associated with *A. bigeloviae,* and 2 exclusive and 15 total species with *A. trixa*. The majority of gall associate species were parasitoid wasps (73.9%), but a predatory beetle, inquiline weevil, hypergalling midge, and hemipteran herbivores were also found. Total abundance was highly variable among the sites and gall morphs, ranging from 95-5540 organisms per site per year.

**Table 1.**
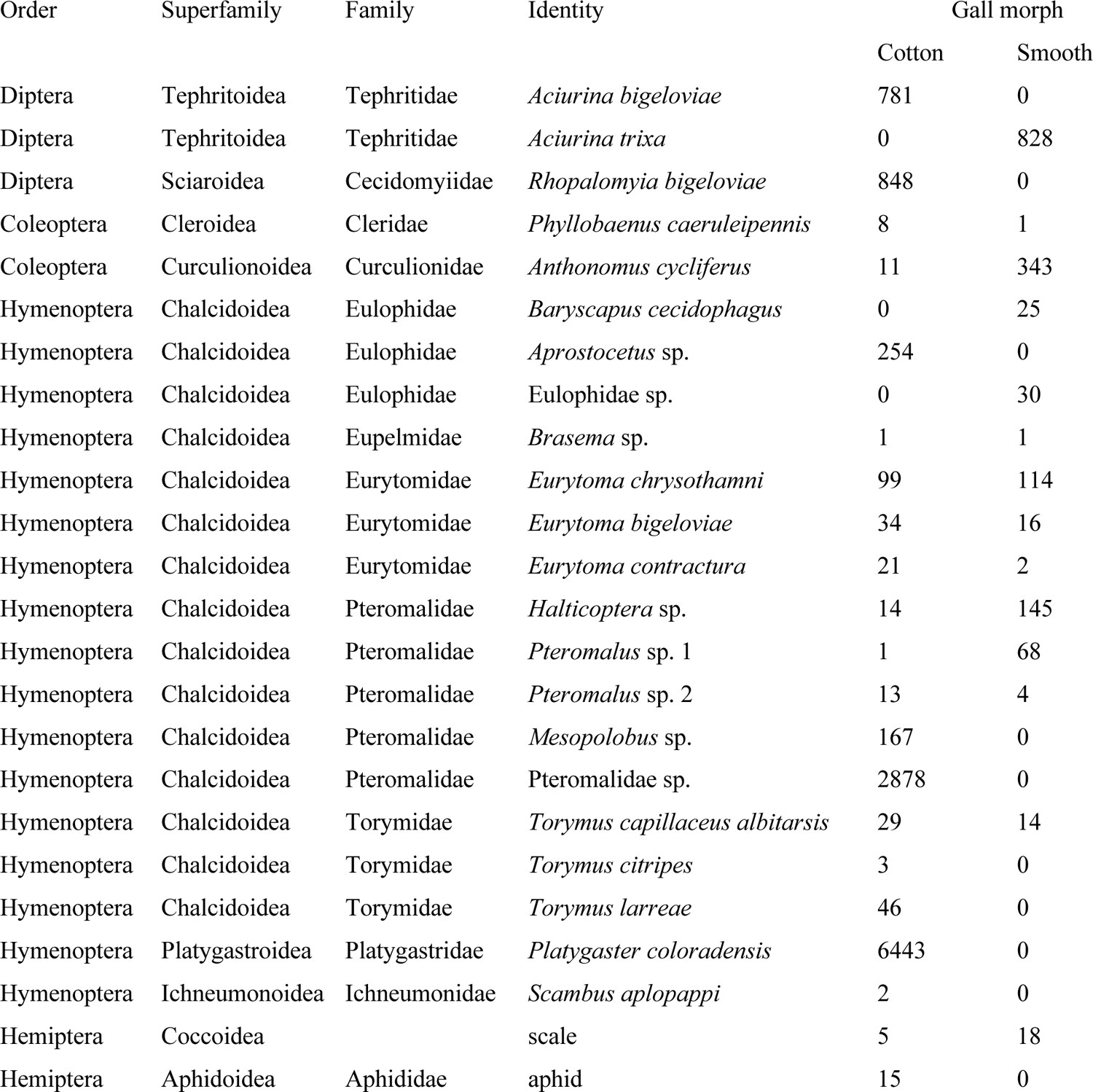
Total counts of individuals of each species identified in this study reared from cotton (*A. bigeloviae* induced) and smooth (*A. trixa* induced) galls.

### Species characterization

*<u>Eurytoma chrysothamni</u>* <u>Bugbee 1975</u> (Fig. 2A-B) (Chalcidoidea: Eurytomidae) was identified to genus using (Gibson et al. 1997). Specimens collected from *Aciurina* galls in 1982 and 1983 by Dodson (Dodson and George 1986) at the MSB were originally identified as *Eurytoma obtusiventris* Gahan 1934, a well-known parasitoid of *Eurosta solidaginis* on goldenrod, which has not been recorded from the southwest. We were able to obtain *E. obtusiventris* specimens (GenBank OR336231) and determined both from morphological examination and from COI sequence (GenBank OR336232, 87.7% similarity) that specimens reared in this study are not conspecific. Species was determined using morphological description (Bugbee 1975) and host association (Wangberg 1981, Bugbee 1982) as no previously generated genetic data was available. The *E. chrysothamni* sequence generated here is most similar (88.41%) to *Eurytoma rhois* (GenBank EF185154). *Eurytoma chrysothamni* was the most abundant parasitoid, was present at all but one site, and was more abundant in smooth galls than cotton galls (115 smooth: 99 cotton). Dissections confirmed *E. chrysothamni* is a koinobiont endoparasitoid.

**Figure 2.**
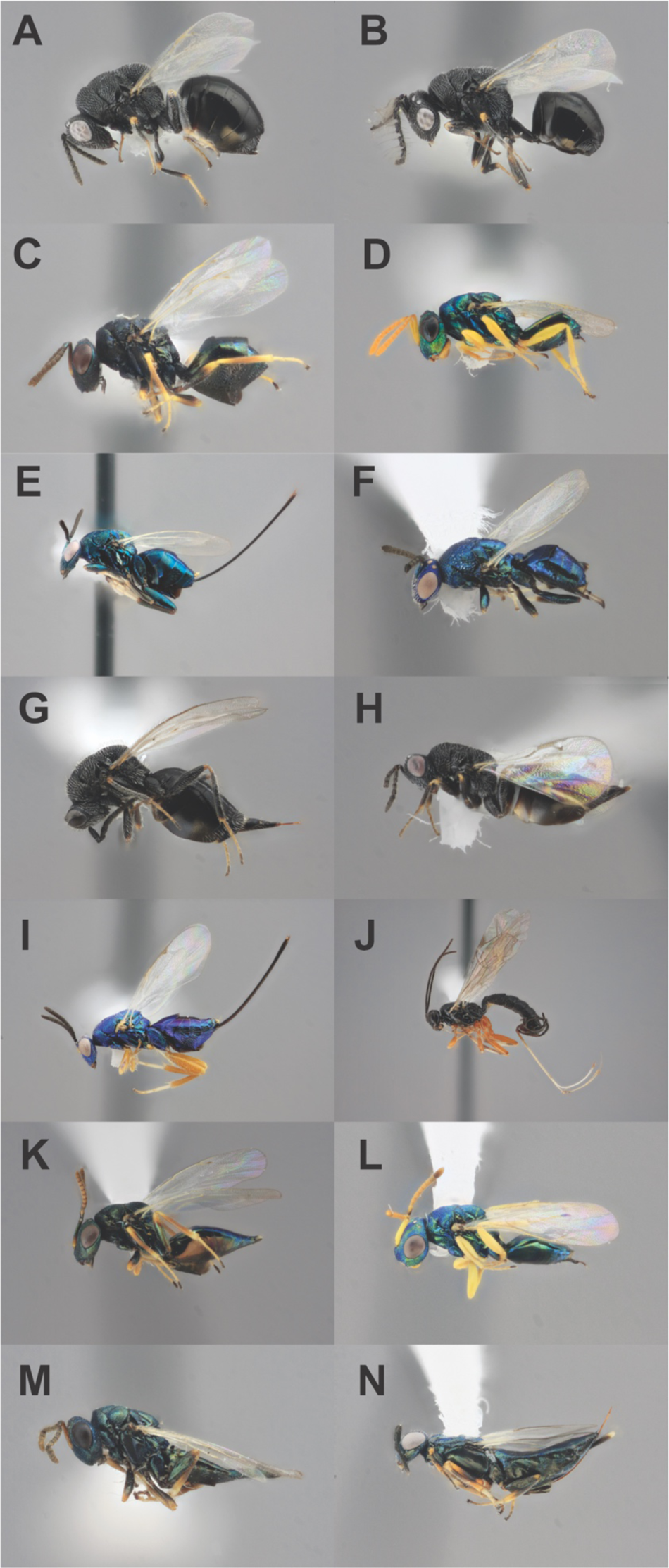
Primary parasitoid representative specimens and MSB catalog numbers. A. *Eurytoma chrysothamni* female MSBA81962. B. *E. chrysothamni* male MSBA81963. C. *Halticoptera* sp. female MSBA81968. D. *Halticoptera* sp. male MSBA81969. E. *Torymus capillaceus albitarsis* female MSBA81960. F. *T. c. albitarsis* male MSBA81961. G. *Eurytoma bigeloviae* female MSBA81964. H. *Eurytoma contractura* female MSBA81965. I. *Torymus citripes* female MSBA81959. J. *Scambus applopappi* female MSBA81958. K. *Pteromalus* sp. 1 female MSBA81966. L. *Pteromalus* sp. 1 male MSBA81967. M. *Pteromalus* sp. 2 female MSBA81970. N. *Brasema* sp. female MSBA81971. Photo credit Q. Baine.

*<u>Eurytoma bigeloviae</u>* <u>Ashmead 1890</u> (Fig. 2G) was similarly identified to genus with (Gibson et al. 1997), and to species using description (Ashmead 1890) and host association (Noyes 2019). Its sequence (GenBank OR336233) was 94.94% similar to *Eurytoma spongiosa* (BOLD PIPI061-09.COI-5P), 86.9% similar to *E. chrysothamni,* and 88.0% similar to *E. obtusiventris.* This species was twice as abundant in the cotton than smooth galls (34:16) and occurred at seven sites. It was present in both morphs from one sympatric site (18 cotton, 4 smooth); however, it was only present on cotton galls at the second sympatric site. *Eurytoma bigeloviae* was unique in phenology, while most other parasitoids emerged in the summer closely following *Aciurina’s* emergence; larvae of *E. bigeloviae* were found feeding within the galls in late May, the first specimen emerged in October and many pupae were still present and alive during late-season dissections in mid-December (Fig. 3). This species is likely thelytokous parthenogenetic as no males were discovered. Dissections and larval sequencing (GenBank OR336224, 100% similarity) confirmed *E. bigeloviae* is an idiobiont ectoparasitoid.

**Figure 3.**
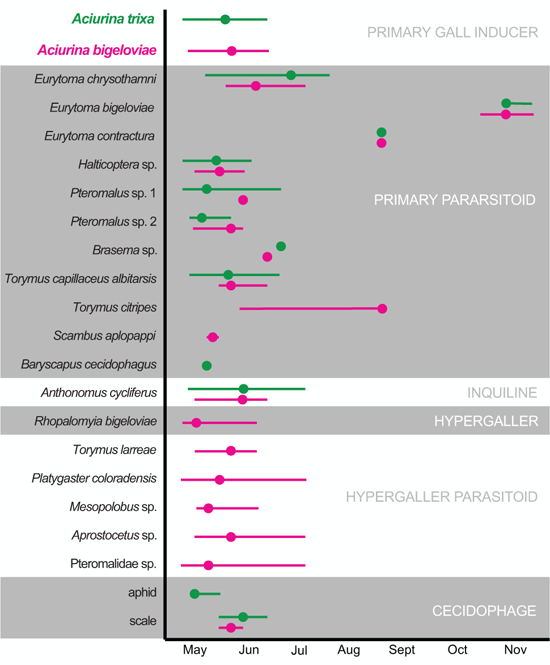
Adult emergence periods, represented as ranges (lines) with means (circles), of each species identified in this study combined from two years of data collection and arranged by ecological guild.

*<u>Eurytoma contractura</u>* <u>Bugbee 1967</u> (Fig. 2H) closely matches description in (Bugbee 1967). The sequence (GenBank OR336223) is 94.65% similar to a private *Eurytoma gigantea* on BOLD, 90.34% similar to *Eurytoma rhois* (GenBank EF577427), 95.6% similar to *E. bigeloviae*, 86.4% similar to *E. chrysothamni*, and 88.4% similar to *E. obtusiventris*. *Eurytoma contractura* was much more abundant in the cotton galls versus the smooth (21:2). Like *E. bigeloviae*, only females of this species were discovered; however, males are reported for this species from the type locality (Bugbee 1967). Though not as late as *E. bigeloviae*, *E. contractura* also emerged later than the other parasitoids, and well beyond the emergence period of the inducer, in August (Fig. 3).

An unidentified *<u>Halticoptera</u>* <u>species</u> (Fig. 2C-D) (Chalcidoidea: Pteromalidae), was identified using (Gibson et al. 1997) and was 99.81% similar in sequence (GenBank OR336230) to an unidentified Pteromalidae (BOLD NGNAI1829-13.COI-5P) from British Columbia of BIN BOLD:ACJ0601. *Halticoptera* sp. was highly abundant overall, but was ∼10x more common in smooth galls (145 smooth: 14 cotton) and present at all smooth sites. It was present in both morphs from one sympatric site (67 smooth: 4 cotton); however, it was only present on smooth galls at the second sympatric site. *Halticoptera* is recorded as an endoparasite on *Aciurina* spp. (Wangberg 1981).

An unidentified *<u>Pteromalus</u>* <u>species</u> (Fig. 2K-L) (Chalcidoidea: Pteromalidae) identified with (Gibson et al. 1997), is 98.54% similar in sequence (GenBank OR438294) to an unidentified Pteromalidae (NGAAB697-14.COI-5P) from British Columbia in BIN BOLD:AAU9358. Like *Halticoptera* sp., *Pteromalus* sp. is far more abundant in smooth galls (68 smooth: 1 cotton), and was most abundant (29 smooth) at the site in Albuquerque, NM. Unlike *Halticoptera* sp., *Pteromalus* sp. 1 was specific to smooth galls at both the sympatric sites (15 smooth: 0 cotton).

A second unidentified *<u>Pteromalus</u>* <u>species</u> (Fig. 2M) was also identified with (Gibson et al. 1997) and is 99.84% similar in sequence (GenBank OR336228) to an unidentified Pteromalidae (GenBank KM564952) from Alberta in BIN BOLD:ACG4086. *Pteromalus* sp. 2 is more abundant in cotton galls (4 smooth: 13 cotton) and was not present at either sympatric site.

An unidentified *<u>Brasema</u>* <u>species</u> (Fig. 2N) (Chalcidoidea: Eupelmidae) was identified with (Gibson et al. 1997); the sequence (GenBank OR438295) is 99.1% similar to *Brasema* “sp. GG2” (BOLD NGNAO1018-14.COI-5P). Only 2 individuals emerged and were counted: 1 from each gall morphology. This likely represents a generalist parasitoid that occurred facultatively on *Aciurina*.

*<u>Torymus capillaceus albitarsis</u>* <u>Huber 1927</u> (Fig. 2E-F) (Chalcidoidea: Torymidae) females were identified to species using (Grissell 1976, Gibson et al. 1997) and males were determined conspecific by 99.4% nucleotide similarity between the COI sequences generated (GenBank OR336229 and OR336225). The female sequence is 99.46% identical to an unidentified Torymidae from California (BOLD BBHYA2339-12.COI-5P). Twice as many specimens were reared from cotton as smooth (29:14). Only a single individual was associated with a smooth gall from a sympatric site (1 smooth: 7 cotton). Larval sequencing and dissections confirmed *T. c. albitarsis* is an idiobiont ectoparasitoid.

*<u>Torymus citripes</u>* <u>Huber 1927</u> (Fig. 2I) were also identified using (Grissell 1976, Gibson et al. 1997). Only 3 specimens total were collected and all were from allopatric cotton galls at two sites. The sequence (GenBank OR336222) is 92.54% similar to *Torymus* sp. (BOLD ROSE614-08.COI-5P) of BIN BOLD:AAG4354. *Torymus citripes* is recorded as an idiobiont ectoparasitoid on other tephritid hosts (Emlen 1992).

*<u>Scambus aplopappi</u>* <u>(Ashmead 1890)</u> (Fig. 2J) (Ichneumonoidea: Ichneumonidae: Pimplinae) was identified with (Goulet and Huber 1993, Khalaim and Ruíz-Cancino 2022). Only two female specimens were collected, both from a single allopatric cotton site in 2022. The sequence (GenBank OR336234) is 98.24% similar to a private *Scambus* sequence on BOLD, and 94.15% to *Scambus vesicarius* (GenBank KT599318), a willow gall sawfly parasitoid. *Scambus aplopappi* is an idiobiont ectoparasitoid (Khalaim and Ruíz-Cancino 2022).

*<u>Baryscapus cecidophagus</u>* <u>(Wangberg 1977)</u> (Chalcidoidea: Eulophidae) is the only identified parasitoid from the primary chamber that was not solitary; 25 individuals emerged from the inside of a single *A. trixa* exuvium from one smooth gall at a sympatric site. The species was identified to family with (Gibson et al. 1997), and to genus by sequence (GenBank OR336226), which is 93.67% similar to *Baryscapus* sp. (BOLD NGNAO1315-14.COI-5P) from British Columbia. Species identification was made from the description and host association (Wangberg 1977, 1981). This gregarious species does not require an elongate ovipositor to reach the central chamber. Because *Aciurina* larvae chew themselves a “trap door” (Fig. 4C) to allow themselves to exit the gall as adults without chewing mouthparts, there is a small section of the gall surface that leads directly to the central chamber. *Baryscapus cecidophagus* takes advantage of this and the adult chews through this thin-walled section to oviposit into the host and complete its life cycle as an endoparasitoid (Wangberg 1977).

**Figure 4.**
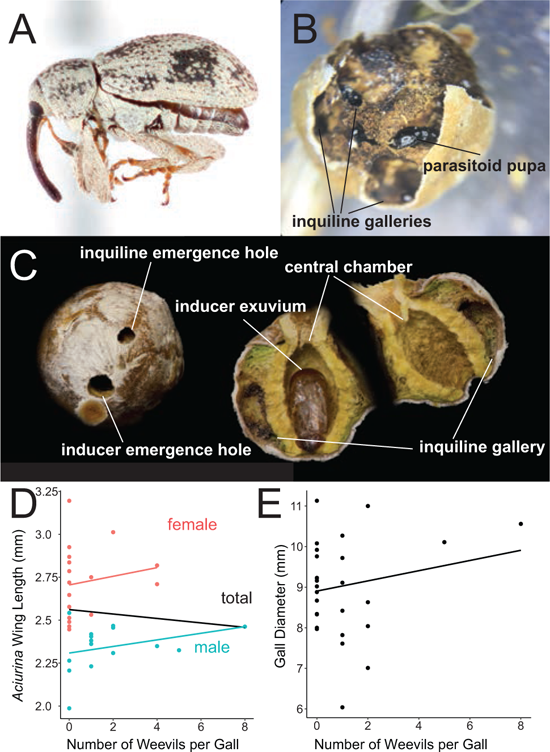
The inquiline *Anthonomus cycliferus* weevil. A. adult habitus. B. Eulophidae sp. parasitoid pupae within weevil galleries. C. *Aciurina trixa* gall exterior and interior with weevil damage. Number of successful weevil emergences compared to: D. The correlation between *A. trixa* inducer size estimated by wing length and number of emerged weevils per gall and E. The correlation between gall diameter and number of emerged weevils per gall. Photo credit Q. Baine.

*<u>Anthonomus cycliferus</u>* <u>Fall 1913</u> (Fig. 4A) (Coleoptera: Circulionidae) were identified using keys in (Clark and Burke 2005). The sequence (GenBank OR336227) obtained aligns to a private *Anthonomus* sp. on BOLD with 98.75% similarity. They bore galleries that fill with frass as they feed, and then chew rough circular holes through the dermal layer to emerge as adults (Fig. 4C). Weevils are obligate inquilines (Wangberg 1981) that feed on the parenchymatous tissue between the outer dermal tissue and sclerenchyma (nutritive tissue consumed by inducer larva) of the inducer chamber, and can be present in high density (maximum count per gall = 8). More than one weevil often emerges from a single gall. When present, the mean adult count was 1.97. The occurrence of weevils did not directly cause inducer mortality: 51.2% of galls with inquiline success also had successful inducer emergence compared to 44.5% inducer success in smooth galls with no weevil emergence. Neither inducer body size (both sexes: *F*=0.26, df=30, *P*=0.611; female: *F*=0.41, df=15, *P*=0.53; male: *F*=1.43, df=13, *P*=0.254) nor gall size (*F*=0.96, df=27, *P*=0.335) were inversely correlated with the number of emerged weevils (Fig. 4D-E), suggesting that the weevils are not siphoning plant-derived resources or otherwise negatively impacting the inducer fitness. Weevils were far more abundant in smooth galls than cotton (343:11), though this difference was weighted heavily by an extreme density from a single site in the Magdalena mountains; 42.8% of galls from this site in 2022 hosted weevils for a total of 255 individuals.

One unidentified <u>Eulophidae</u> (Chalcidoidea) species pupae was discovered during late-season dissections, specifically within the larval galleries of the inquiline weevil (Fig. 4B). Sequencing confirmed that this species is distinct from *Aprostocetus* sp., which parasitizes the hypergaller midge (see below), and *Baryscapus cecidophagus*, which parasitizes the inducer, indicating host specificity to the weevil. The sequence (GenBank OR438293) was 97.87% similar to an unidentified Eulophidae (BOLD MPGK1929-19.COI-5P) collected in Missoula, Montana. The eulophid may be a generalist or semi-specialist parasitoid that is facultative on the weevil because it resides closer to the gall surface and is therefore easier to access.

*Phyllobaenus caeruleipennis* (Wolcott 1908) (Coleoptera: Cleridae) was a predatory beetle identified using (Opitz 2010, Leavengood Jr 2014). *Phyllobaenus caeruleipennis* only occurred at three sites, and was more common in cotton galls (8 cotton, 1 smooth). The genus is cited as generally predatory on gall inducers, and is recorded in association with tephritid flies (Wangberg 1980).

The hypergaller midge *Rhopalomyia bigeloviae* (Cockerell 1889) induces a novel form of secondary gall on the surface of the cotton *A. bigeloviae* galls, but are not found on the smooth *A. trixa* galls in New Mexico, though they have been previously recorded on *A. trixa* galls in California (Gagné 1989, Russo 2021). The midges have their own specific suite of 5 species of hymenopteran parasitoids (*Torymus larreae* Grisell 1976, *Platygaster coloradensis* Ashmead 1893, *Mesopolobus* sp., *Aprostocetus* sp., and Pteromalidae sp.), which in some cases are present in extremely high abundances. This sub-community and its impact on *A. bigeloviae* ecology are fully described in (Baine et al. 2023).

Some herbivorous hemipterans (aphids and scale insects) were collected as gall-associates because they were not visible at time of collection but are likely generalist herbivores on plant tissue. Two sets of predators were not included in the analyses of this dataset: 1) a set of reared <2 mm in length hymenopterans whose association with the gall, as opposed to a section of stem, in particular could not be confirmed, and 2) an unidentified bird species, which heavily predated cotton galls at one collection site; however, these predated galls were not collected as part of this study.

## Discussion

This study is the first to fully characterize the entire arthropod communities associated with *A. bigeloviae* and *A. trixa* and found that both are rich and trophically complex in New Mexico. The aggregate community includes 24 species and at least 6 guilds: 2 gall inducers, a predator, 2 inquilines (including a hypergall inducer), 6 inquiline parasitoids, 11 primary parasitoids and 2 cecidophages.

Our community characterizations are supported by previous partial studies on *Aciurina* associates. *Anthonomus* inquiline weevils and *Eurytoma chrysothamni* have been reported in *A. trixa* smooth galls by Wangberg (Wangberg 1981) in Idaho, and the genera *Torymus, Eurytoma, Halticoptera*, and *Platygaster* are associated with several *Aciurina* species in the west (Wangberg 1981, Headrick et al. 1997, Noyes 2019). *Rhopalomyia* has also been recorded in both *Aciurina* systems (Gagné 1989, Russo 2021). Incidental unpublished parasitoid collections from work by Dodson on *Aciurina* in New Mexico (Dodson and George 1986, Dodson 1991) housed at the MSB also include some of these genera. Surprisingly, only a single community has been partially characterized for *A. trixa* in Idaho (Wangberg 1981) and no communities have been formally characterized for *A. bigeloviae.* The *A. trixa* partial survey included 17 morphospecies, including an obligate inquiline weevil, “secondary gall-former” midge and 10 parasitoid wasps. At least 9 of the listed associates (Wangberg 1981) were not identified in the current study, so there may be local diversity present among different *A. trixa* populations.

Both of these *Aciurina*-associated communities appear to be far richer than those documented from other studied Asteraceae-galling Tephritidae species (e.g. *Valentibulla* spp., *Tephritis conura, Eutreta diana, Eutreta xanthochaeta, Tomoplagia rudolfi, Urophora cardui*) by at least two-fold (Wangberg 1978, Romstock-Volkl 1990, Emlen 1992, Hernández-López et al. 2021, Maia and Silva 2021, Swenson et al. 2022). The most closely-related tephritid fly with a species-level community description, *Eurosta solidaginis*, only hosts 3 associates (Walton 1988). Furthermore, we find that the *Aciurina bigeloviae* community in particular is the most guild rich and most complex trophic web so far described among this group; there are not only associated herbivores, predators and primary parasitoids, but also 2 obligate inquilines each with their own parasitoid communities. The genus *Aciurina* includes several other species that all occur in western North America on at least 4 host Asteraceae species, and many can be highly abundant in their respective ecosystems (Wangberg 1981, Steyskal 1984, Russo 2021). Additional characterizations of these species could reveal even more undescribed richness and support our understanding of the drivers of biodiversity.

While their associations with *Aciurina* are now known, the full ecologies/life histories of many of these gall associates still have gaps. Most of the primary parasitoids identified in this meta-community emerge as adults in the summer, which presents a mystery because the *Aciurina* gall is not visible until fall, 3-5 months after oviposition. However, if we combine known natural history with our functional guild assignments, we are able to make some educated guesses about the ecology of these associates: 1) *Torymus capillaceus albitarsis* and *T. citripes* are both described in association with other gall midge or tephritid hosts on *Ericameria* and *Artemisia* (sagebrush) (Grissell 1976, Emlen 1992, Noyes 2019), therefore it seems likely that these species represent generalists that find alternate hosts in the summer season. Because *Torymus* have elongated ovipositors, and at least one is an idiobiont ectoparasitoid, we presume that they attack *Aciurina* galls in the fall when they are full sized. 2) Both *Halticoptera* and *Pteromalus* spp. are broadly gall-associated parasitoid genera, so these unidentified species may be generalists with alternate hosts, similar to *Torymus*. However, because *Halticoptera* sp. has a short ovipositor and is an endoparasitoid (Wangberg 1981), it may instead have a life history similar to that of *E. chrysothamni* and attack *Aciurina* prior to gall growth. Barcode sequencing of *Aciurina* early-instar larvae and other *Ericameria* and *Artemisia* gall inducers may reveal oviposition timing of these parasitoid species and confirm these relationships in future studies.

The importance of identifying the impact of key ecosystem engineers in maintaining threatened biodiversity is often overlooked (Boogert et al. 2006), particularly in insects, where large-scale species loss (Rhodes 2019, Raven and Wagner 2021) may be in danger of outpacing species description (Foottit and Adler 2009, Audisio 2017). Though rich and widespread, with an estimated 21,000 species worldwide (Takeda et al. 2021), gall-inducing insects are under-described and their numerous interactions and associates are largely undocumented (Redfern 2011, Takeda et al. 2021, Ward et al. 2022). Because of their essential role in sustaining ecological diversity, ecosystem engineers and the communities they support require description to provide accurate estimates that advance species conservation (Boogert et al. 2006, Bouma et al. 2009). Furthermore, the strength of ecosystem engineers in community stabilization requires experimental investigation to inform responsible ecological management in the face of changing environments (Boogert et al. 2006, Mullan Crain and Bertness 2006, Bouma et al. 2009). As climate change and agricultural expansion continue to decimate insect populations (Rhodes 2019), the species and ecosystem services dependent on gall inducers may be lost to the world before they can be protected (Foottit and Adler 2009, Audisio 2017), resulting in cascading trophic effects that could impact pest control, invasive species, pollination, soil health and biodiversity (Bouma et al. 2009, Rhodes 2019, Snyder 2019).

## Acknowledgments

We thank D. Lightfoot and K. Miller for use of MSB Division of Arthropods mounting and photography resources; James Winston Cornish for beetle identification; C. Valderrama Hincapié for Spanish language editing, and C. Valderrama Hincapié, I. Gulisija-Radic, J. Booker, V. Wilson, E. Carabotta, A. Brackeen, M.K. Poling, P. Zedalis, and M. Keller for help with gall collection and rearing. Research and collections were carried out on land historically unceded by Tiwa, Pueblos, Piro, Zuni, Apache, Ute, and Diné people. This work was supported in part by UNM Biology Graduate Student Association GRAC and Graduate and Professional Student Association grants in 2021 and 2022.

